# Systematic whole-genome sequencing reveals an unexpected diversity among actinomycetoma pathogens and provides insights into their antibacterial susceptibilities

**DOI:** 10.1101/2022.01.03.474876

**Authors:** Andrew Keith Watson, Bernhard Kepplinger, Sahar Mubarak Bakhiet, Nagwa Adam Mhmoud, Michael Goodfellow, Ahmed Hassan Fahal, Jeff Errington

## Abstract

Mycetoma is a neglected tropical chronic granulomatous inflammatory disease of the skin and subcutaneous tissues. More than 70 species with a broad taxonomic diversity have been implicated as agents of mycetoma. Understanding the full range of causative organisms and their antibiotic sensitivity profiles are essential for the appropriate treatment of infections. The present study focuses on the analysis of full genome sequences and antibiotic resistance profiles of actinomycetoma strains from patients seen at the Mycetoma Research Centre in Sudan with a view to developing rapid diagnostic tests. Seventeen pathogenic isolates obtained by surgical biopsies were sequenced using MinION and Illumina methods, and their antibiotic resistance profiles determined. The results highlight an unexpected diversity of actinomycetoma causing pathogens, including three *Streptomyces* isolates assigned to species not previously associated with human actinomycetoma and one new *Streptomyces* species. Thus, current approaches for clinical and histopathological classification of mycetoma may need to be updated. The standard treatment for actinomycetoma is a combination of sulfamethoxazole/trimethoprim and amoxicillin/clavulanic acid. Most tested isolates were not susceptible to sulfamethoxazole/trimethoprim or to amoxicillin alone. However, the addition of the β-lactamase inhibitor clavulanic acid to amoxicillin increased susceptibility, particularly for *Streptomyces somaliensis* and *Streptomyces sudanensis. Actinomadura madurae* isolates appear to be particularly resistant under laboratory conditions, suggesting that alternative agents, such as amikacin, should be considered for more effective treatment. The results obtained will inform future diagnostic methods for the identification of actinomycetoma and treatment.

**Author Summary:** Mycetoma is a common health and medical problem that is endemic in many tropical and subtropical countries and has devastating effects on patients. The destructive nature of late-stage infection means that treatment often requires long term use of antibiotic therapy, massive surgical excisions and amputation. Several different bacterial species have been described as causing this disease but our understanding of the true diversity of mycetoma causing bacteria has been limited by a lack of molecular sequence data. We have now sequenced the genomes of 17 samples isolated from patients at the Mycetoma Research Centre in Sudan, revealing a diverse range of species associated with infection including one new *Streptomyces* species, and three species with no previous association with human mycetoma. Crucially, all isolates had a high level of resistance against the current first-line antibiotics used to treat actinomycetoma under laboratory conditions. This resistance was strongest in *Actinomadura madurae*, which was also the most frequently observed species isolated from patients in our study. We hope that these results will aid in the development of future rapid diagnostic tools and the improvement of treatment outcomes.

## Introduction

Mycetoma, a neglected tropical disease, is a chronic subcutaneous granulomatous inflammatory disease [1,2]. The inflammatory process usually spreads to affect the skin, deep tissues and bone, leading to massive destruction, deformity and disability, and can be fatal [3–5]. It is endemic in many countries around the equator in a zone often described as the mycetoma belt, with Sudan reported as the most affected country [6–8].

Mycetoma is characterised by a painless mass with multiple sinuses which discharge material including grains which are colonies of the causal agents. Colours, sizes and consistency of the grains can often be indicative of the aetiological agent [9,10]. It can affect different body parts but is most commonly seen in the feet and hands. Young adults and children are frequently affected, leading to numerous negative medical, health and socioeconomic impacts on patients and their families [5,11]. The combination of the painless nature of the initial stages of the disease, low availability of health facilities in endemic regions and the low socioeconomic status of those infected explains the late presentation of patients at clinics. Consequently, extensive surgical intervention often becomes the only remaining option for treatment [12–14].

While mycetoma infections are characterised by a shared phenotype, a broad taxonomic range of species have been described as causative agents, including fungi producing eumycetoma and actinobacteria causing actinomycetoma [6,15,16]. The most common agents of actinomycetoma are *Actinomadura madurae, Actinomadura pelletieri, Nocardia asteroides, Nocardia brasiliensis* and *Streptomyces somaliensis [17]*; less well known species include *Actinomadura latina [18,19] Actinomadura mexicana* [20], *Streptomyces albus* [21], *Streptomyces griseus* [22] and *Streptomyces sudanensis* [23]. Historically, mycetoma diagnosis has been based mainly on the clinical presentation, surgical biopsies and histopathological examination of the grains in the mycetoma granuloma mass and on culture media, but only in a few centres [24–26]. Unfortunately, these features often overlap between different causative organisms, making diagnosis challenging [27,28]. There is also evidence that streptomycetes can cause mycetoma in animals [29,30], as well as fistulous withers in donkeys [31].

Accurate diagnosis and the ability to distinguish between actinomycetoma and eumycetoma is vital, as the treatments for them are fundamentally different [24–26]. Current treatment for eumycetoma involves long term anti-fungal medication and surgical intervention. On the other hand, actinomycetoma is treated with a combination of antibiotics. However, prolonged treatment is required to effect a cure, and the response to treatment is variable depending on the causative agents and associated drug resistance [32–35].

It is important to identify the causative agents of actinomycetoma to ensure they respond to medical treatment. Since actinomycetoma is caused by several species, it is possible that molecular differentiation within and between species may reveal differences in responses to frontline treatments and that adaptation of treatments would be beneficial for patient care. We, therefore, set out to obtain whole-genome sequences of a variety of actinomycetoma pathogens, determined their taxonomic status, and characterised their resistant profiles to commonly used antibiotics.

## Materials and Methods

### Sample isolation and subculture

The patients were seen at the Mycetoma Research Center (MRC), University of Khartoum. Granulomatous material from mycetoma lesions were obtained by surgical biopsy. The grains were washed three times in sterile normal saline, then plated on yeast extract agar (YEA) and incubated at 37°C for one to two weeks under aerobic conditions. Once microbial growth was visible, the colonial morphology was reported, and Gram-stained smears were prepared from each isolate. Actinomycetoma organisms, unlike their eumycetoma counterparts, are characterised by their typical filamentous Gram-positive appearance.

Strains received at Newcastle University from the MRC were cultivated on tryptic soy agar and oatmeal agar (20 g/l of oat was boiled in water for 20 min; the liquid was strained using a sieve, and 20 g/l of agar was added prior to autoclaving) at 30°C.

### Methods for histopathology and cytology

Deep excisional biopsies were taken from the mycetoma lesions and preserved in 10% neutral buffer formalin. The tissue biopsies were processed and then cut using a rotary microtome (Leica, Germany). The 3-5μm sections obtained were stained with haematoxylin and eosin (H&E), and the slides examined using a light microscope (Olympus, Germany) for the presence of grains and the type of inflammatory reactions according to previously described criteria [26].

The cytological smears were prepared on aspirated materials from mycetoma lesions using a 23 gauge needle attached to a 10 ml syringe. The smears were allowed to air dry and then stained with May-Grünwald-Giemsa (MGG) and H&E stains. The slides were then examined by light microscopy for the presence of grains of the causal agents using previously described criteria [36].

## Ethics statement

Ethics approval for this study was obtained from the Mycetoma Research Centre, Khartoum, Sudan IRB (Approval no. SUH 11/12/2018). Written informed consent was obtained from each adult patient and parents or guardians of the population under 18 years old. Confirmed mycetoma cases were referred for management at the Mycetoma Research Centre (MRC).

### DNA isolation, sequencing and assembly

DNA was isolated using a modified version of the “salting-out” method [37]. A 15 mL culture grown in TSB was harvested by centrifugation and resuspended in 5ml of SET buffer, containing a final concentration of 1.5 mg/ml lysozyme, and incubated at 37°C for 1.5 h. RNase was added to a final concentration of 20 μg/ml and the sample incubated at room temperature for 1 min. Pronase (final concentration 0.5 mg/ml) and SDS (final concentration 1%) were added and mixed by inversion at 37°C for 2 h. 2 ml of 5M NaCl and 5 mL chloroform were added, with incubation at room temperature for 30 min. Phases were separated by centrifugation, and the aqueous phase transferred into a fresh tube. The DNA was precipitated by the addition of 0.6 vol of propan-2-ol. The high molecular weight DNA was spooled around a sealed Pasteur pipette. The DNA was washed using freshly prepared 70% ethanol before being allowed to air dry. The DNA was finally dissolved in 10 mM Tris-HCl pH 8.5.

The libraries for Minion sequencing were prepared using the ligation sequencing kit with the native barcode extension kit, according to the protocols of the manufacturer. Sequencing was performed on two Flo-MinSP6 flow cells. Libraries for Illumina sequencing were prepared using the Illumina DNA prep kit with Nextera™ DNA CD Indexes. Libraries were sequenced on an ISeq 100.

Nanopore reads were base-called and demultiplexed using guppy 3.4.4+a296acb. Draft genome assemblies from nanopore sequences were generated using flye 2.8.2 [38], followed by consensus correction using Medaka 1.2.1 [39] implemented in the “demovo” pipeline (kindly provided by Demuris Ltd). Illumina reads were mapped to the draft Nanopore assembly using minimap 2.17, and pilon 1.23 was used to correct the genome assembly based on the Illumina mapping. This process was repeated four times. Diamond and MEGAN were used to correct frameshifts in the genome assemblies as described previously [40]. Finally, genome assemblies were annotated using the NCBI pgap pipeline [41].

### 16S rRNA taxonomy

The longest 16S rRNA annotated in each isolate genome was used as a query in the EzBioCloud 16S rRNA database. Isolates were assigned to the genus “*Actinomadura*” or “*Streptomyces*” based on their best hits in this database as determined by sequence identity. The 50 most similar rRNA sequences to each isolate sequence were extracted from the database, pooled according to the predicted genus of the corresponding isolates, and dereplicated. This produced two datasets of 16S rRNA’s with similarity to *Actinomadura* and *Streptomyces* related isolates. Additionally, all *Streptomyces* 16S rRNA sequences generated from a recent survey of soil samples in Sudan and 11 unpublished samples (NCBI access.: EU544231.1 to EU544241), isolated from humans and animals in Sudan, were included in the *Streptomyces* dataset [42].

16S rRNA sequences were aligned using mafft7.453 (--auto mode) [43], and the alignment was trimmed using trimal 1.4 (--automated1) [44]. Phylogenetic trees were inferred with iqtree2 using the best scoring GTR model designated by iqtree’s automated model selection [45]. All phylogenetic trees were midpoint rooted and visualised using iToL [46].

### Whole genome taxonomy

GTDB-Tk [47] was used to classify genomes based on whole-genome data using two methods: maximum-likelihood based placement of genomes into a reference phylogeny based on the bac120 set of 120 marker genes using pplacer [48] and by comparing the average nucleotide identity (ANI) of isolate genomes to those in GTDB [49] using FastANI [50].

The genomes of all “close relatives” of isolates identified by GTDB-tk were extracted from the database. Further, the protein sequences of genes from the GTDB bac120 set of phylogenetic markers [49] were extracted from all isolates, “close relative” GTDB genomes, and type strain genomes. As with 16S rRNA, the dataset was separated into *Actinomadura* and *Streptomyces* related isolates. All markers identified as a single copy in isolate genomes were retained for phylogenetic analysis. These marker sequences were individually aligned using mafft (--auto mode) and trimmed using BMGE (BLOSUM30 model) [51]. Phylogenetic trees based on the concatenation of the trimmed alignments were inferred using iqtree2 [45] using the C20 model [52]. All phylogenetic trees were midpoint rooted and visualised using iToL [46].

FastANI 1.32 [50] was used to estimate the Average Nucleotide Identity (ANI) of all pairs of isolated genomes, reference genomes from type strains of known causes of human actinomycetoma, and the genomes of close relatives to the isolates identified by GTDB-tk, thereby providing a measure of their diversity at the whole genome level. Average Amino acid Identity (AAI) values were calculated between isolates and any close relatives from GTDB with an ANI >90% using EzAAI [53]. Finally, DDH values between isolates and the single genome from GTDB identified by all previous methods as their closest relatives were estimated using GGDC [54]. ANI >95% [55], AAI >95% [56] and DDH > 70% [57] are the typical values used to delineate species boundaries.

### Antimicrobial susceptibility assay

The bacterial strains were plated on Mueller-Hinton agar (Oxoid) except for MRC003, MRC013 and MRC019, which grew poorly on this medium and so were plated onto TSA (Difco). MRC008 did not grow as a lawn under any of the conditions tested and was excluded from the analyses. Antimicrobial susceptibility disks (Oxoid) were loaded with the following compound concentrations: amikacin (30 μg), amoxicillin (25 μg), amoxicillin/clavulanic acid (30 μg), erythromycin (15 μg), gentamicin (30 μg), rifampicin (5 μg) and trimethoprim/sulfamethoxazole 1:19 (25 μg). The disks were placed on the plates immediately after inoculation and incubated at 37°C for five days before measuring zones of inhibition in mm. Sequence-based predictions of antimicrobial resistance profiles were made using AMRfinderPlus [58].

## Results

### Isolation of actinomycetoma pathogens

The grains for culture were collected from confirmed mycetoma patients seen at the Mycetoma Research Centre, University of Khartoum. The patients were all from Sudan, except for one patient from Yemen. All patients had meticulous clinical interviews and examinations after giving written consent. In addition, all of them had surgical biopsies taken under local or general anaesthesia. Microbial cultures obtained from grains were Gram-stained to distinguish actinobacterial from fungal pathogens. Seventeen independently isolated strains identified as filamentous actinobacteria were sent to Newcastle University for further analysis by whole-genome sequencing. All whole genome sequences are deposited on NCBI under BioProject ID PRJNA782605.

### 16S rRNA analyses reveal a high diversity amongst clinical isolates

The initial taxonomic assignment of isolates based on 16S rRNA sequence similarity supported their annotation as actinomycetes, with seven isolates assigned to the genus *Actinomadura* and 10 to the *Streptomyces* genus. Of the *Streptomyces* isolates, the 16S rRNAs of 7 of them were most similar to known causes of actinomycetoma; *Streptomyces sudanensis* (3 isolates; MRC006, MRC007 and MRC017) and *Streptomyces somaliensis* (4 isolates; MRC001, MRC009, MRC013, MRC016). These sequences also formed strongly supported clades with reference rRNA sequences in the phylogenetic analyses (Fig 1A & 1C). The initial taxonomic assignment of three isolates from the genus *Streptomyces*, MRC002, MRC003 and MRC012, did not correspond to previously identified sources of human actinomycetoma, with each branching separately in the 16S rRNA tree (Fig 1B, 1D & 1E), suggesting they may represent three independent new sources of human actinomycetoma from within the genus *Streptomyces*. These relationships are discussed in more detail in light of the whole genome data considered below. Interestingly, the clades including the potentially novel MRC012 and the *S. sudanensis* strain also included 16S rRNA sequences from strains isolated by Sengupta, Goodfellow and Hamid (unpublished) data from lesions of donkeys with fistulous withers in Sudan (Fig 1C & 1E; NCBI accessions: EU544241.1, EU544239.1), suggesting that these strains may be capable of infecting multiple hosts thereby raising the possibility of cross-species transmission of actinomycetoma.

**Fig 1.**
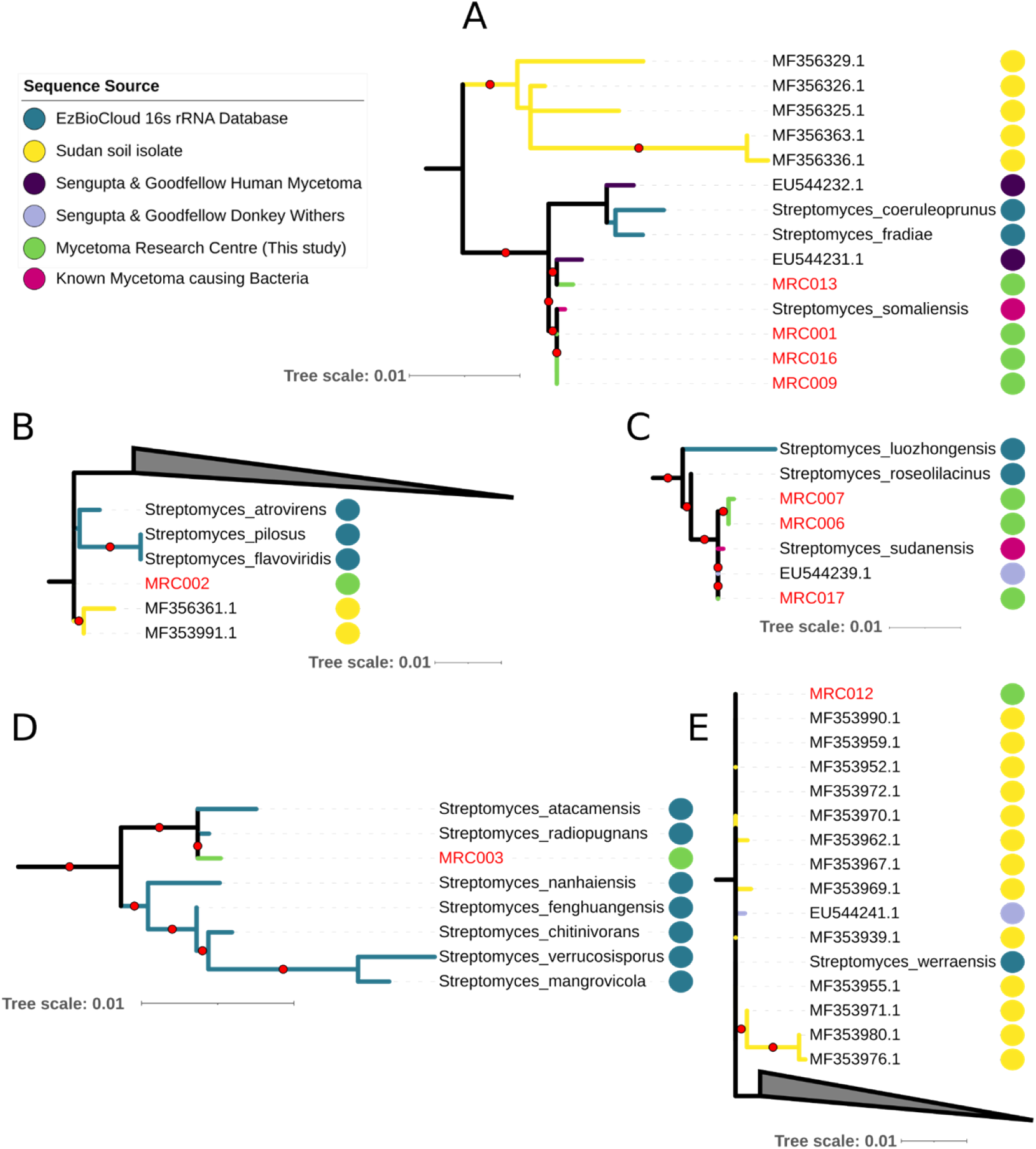
Extracts from the 16S rRNA phylogenies of *Streptomyces* (full figure in SI Fig 1 and https://itol.embl.de/shared/1MX60mtB0Ohk3) showing the placement of isolates (green dots) with related rRNA sequences from the ezBioCloud 16S rRNA database (blue dot), soil isolates from Sudan (yellow dots [42]) and isolates collected by Sengupta, Goodfellow and Hamid (unpublished) from human actinomycetoma (purple) and from donkey withers (lilac). Inferred using iqtree2 and the GTR+F+R5 model. A red dot on branches indicates ultrafast bootstrap support >95.

Six of the seven isolates assigned to the genus *Actinomadura* were from various regions across Sudan; the remaining one was from Yemen (MRC019). The 16S rRNAs of six of the isolates were most similar to the *A. mexicana* 16S rRNA sequence in the EzBioCloud database (>99% identity); the remaining one was closest to the reference *A. madurae* 16S rRNA. In the 16S rRNA phylogeny, all 7 isolates assigned to *Actinomadura* formed a weakly supported clade with *A. madurae* (a known cause of human mycetoma) and *Actinomadrua darangshiensis* (Fig 2), with MRC005 and MRC008 branching at the base of the clade. However, the weak support for this clade and its presence at the foot of the *Actinomadura* tree, means that a relationship between the isolates and *A. mexicana* cannot be excluded. The precise nature of the relationships between these isolates, as well as all other isolates in our dataset, and existing strains were disentangled using corresponding whole genome sequence data.

**Fig 2.**
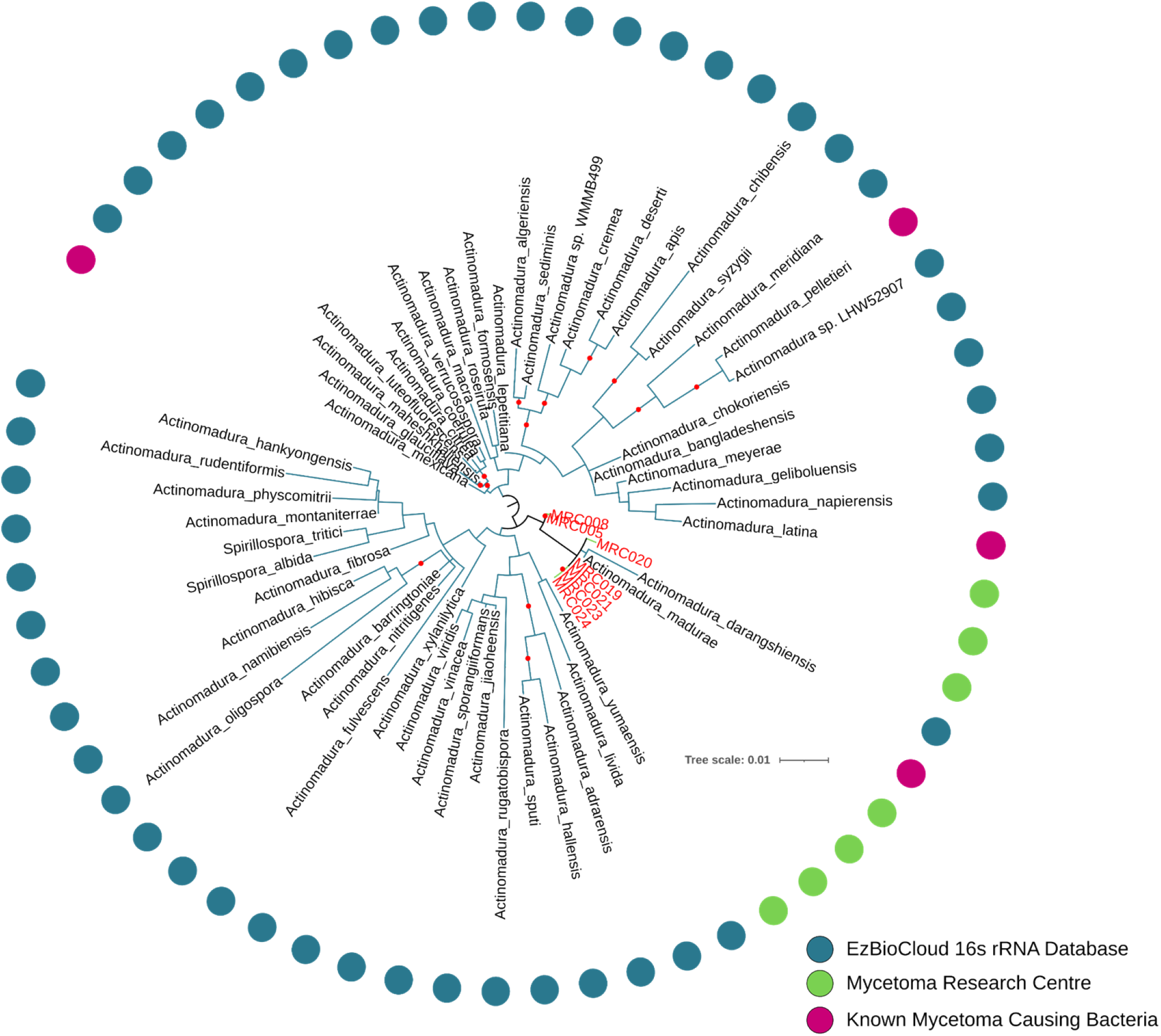
The 16S rRNA phylogeny of *Actinomadura* showing the placement of isolates (green) with related rRNA sequences from the ezBioCloud database (blue; pink for known pathogens). Inferred using iqtree2 and the GTR+F+R4 model. A black dot on branches indicates ultrafast bootstrap support >95. All isolates form a clade with *Actinomadura madurae* and *Actinomadura darangshiensis*, though support for this grouping is low.

### Whole genome analyses support the classification of *S. somaliensis* and *S. sudanensis* as separate species

In 2008, *S. sudanensis* was proposed as a new species closely related to *S. somaliensis* based on a combination of 16S rRNA sequence, DNA:DNA relatedness and phenotypic data *[23]*. Within the Genome Taxonomy DataBase (GTDB), two genomes are annotated as deriving from *S. somaliensis*: GCF_012396115.1 and GCF_000258595.1. However, our comparisons of these genomes to the type strain genomes of *S. somaliensis* and *S. sudanensis,* based on a combination of ANI, AAI and DDH values, revealed that GCF_000258595.1 shares high similarity values to the *S. sudanensis* type strain, which is consistent with its assignment to that species (Table 1). Furthermore, comparison of these genomes provided additional evidence that *S. somaliensis* and *S. sudanensis* should be considered as separate species [23], as ANI, AAI and DDH values are below the thresholds typically used to assign closely related strains to the same species (Table 1).

**Table 1:** Summarising the pairwise ANI, AAI and estimated DDH values of the *S. somaliensis and S. sudanensis* type strain genomes compared to genomes in GTDB. Species thresholds are usually defined by ANI >95%, AAI >95% and DDH >70%. Values that fall below these thresholds are highlighted in grey. Results indicate that GTDB GCF_000258695 (annotated as *S. somaliensis*) is most closely related to *S. sudanensis*, though it is a different species..

| Genome 1 | Genome 2 | ANI (%) | AAI (%) | DDH (GGDH) |
| --- | --- | --- | --- | --- |
| GTDB *S. somaliensis* (GCF_012396115) | *S. somaliensis* Type Strain Genome | 99.6 | 99.7 | 98.4 |
| GTDB *S. somaliensis* (GCF_012396115) | *S. sudanensis* Type Strain Genome | 92.0 | 90.2 | 43.3 |
| GTDB *S. somaliensis* (GCF_000258595) | *S. somaliensis* Type Strain Genome | 91.9 | 89.9 | 42.2 |
| GTDB *S. somaliensis* (GCF_000258595) | *S. sudanensis* Type Strain Genome | 99.8 | 99.9 | 98.5 |
| GTDB *S. somaliensis* (GCF_000258595) | GTDB *S. somaliensis* (GCF_012396115) | 91.79 | 90.1 | 43.3 |

We used GTDB-tk to provide an initial taxonomic assignment of isolates identified by 16S rRNA analysis as related to *S. somaliensis* or *S. sudanensis*, and to identify all close relatives of isolates with publicly available genomes deposited in the GTDB (Table S1). We used these data to construct a multimarker phylogeny for isolates and their close relatives, and for pairwise comparisons of ANI, AAI and DDH similarities. The data from whole genome comparisons between the isolates and the *S. somaliensis* and *S. sudanensis* strains were consistent with the results from the 16S rRNA analyses. All three isolates assigned to *S. sudanensis* in the 16S rRNA phylogeny were found to be most similar to *S. sudanensis* (GCF_000258595.1) by GTDB-tk and formed a strongly supported clade with the *S. sudanensis* type strain and genome GCF_000258595.1 in the multi-marker phylogeny (Fig 3). All ANI, AAI and DDH values between these isolates and the *S. sudanensis* genome GCF_000258595.1 were consistent with the isolate belonging to the same species as the type strain (Table 2; Fig 4; Fig 5). Similarly, all four isolates assigned to *S. somaliensis* in the 16S rRNA gene sequence analysis formed a strongly supported lineage with *S. somaliensis* in the multi-marker tree of the group (Fig 3). In addition, three of the four isolates (the exception MRC013; is discussed below) were also assigned to *S. somaliensis* by GTDB-tk; the ANI, AAI and DDH values between these isolates and the type strains of *S. somaliensis* supported their assignment to this species (Table 2; Fig 4; Fig 5).

**Fig 3.**
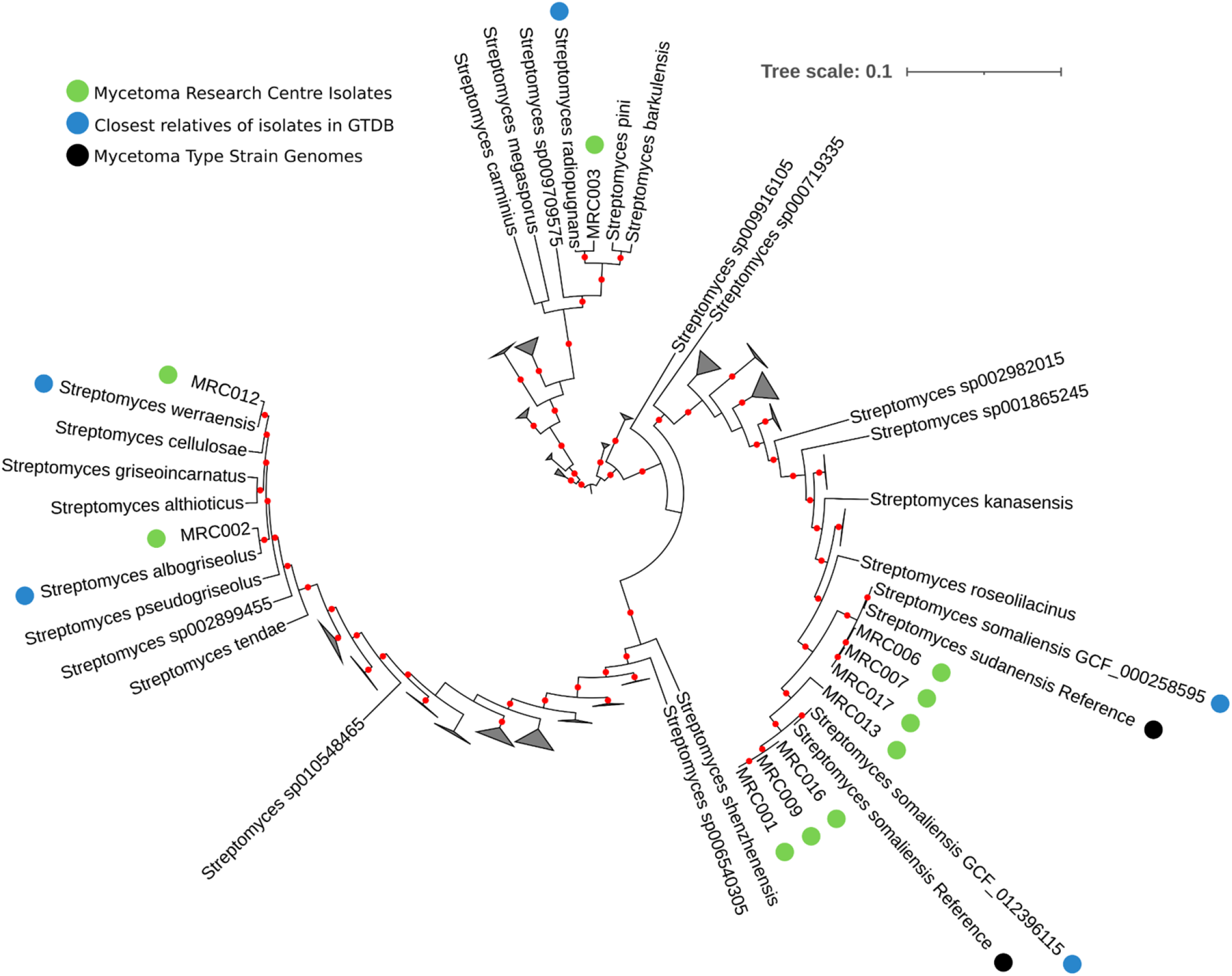
Tree of single copy orthologs belonging to the GTDB bac120 dataset, from the genomes of *Streptomyces* related isolates from the Mycetoma Research Centre (green), their relatives according to ANI in GTDB (with the closest relatives indicated in blue) and genomes from type strains of species typically associated with actinomycetoma (black). A red dot on branches indicates ultrafast bootstrap support >95. The tree was inferred in iqtree2 using the C20 model. The full figure is available at https://itol.embl.de/shared/1MX60mtB0Ohk3.

**Fig 4.**
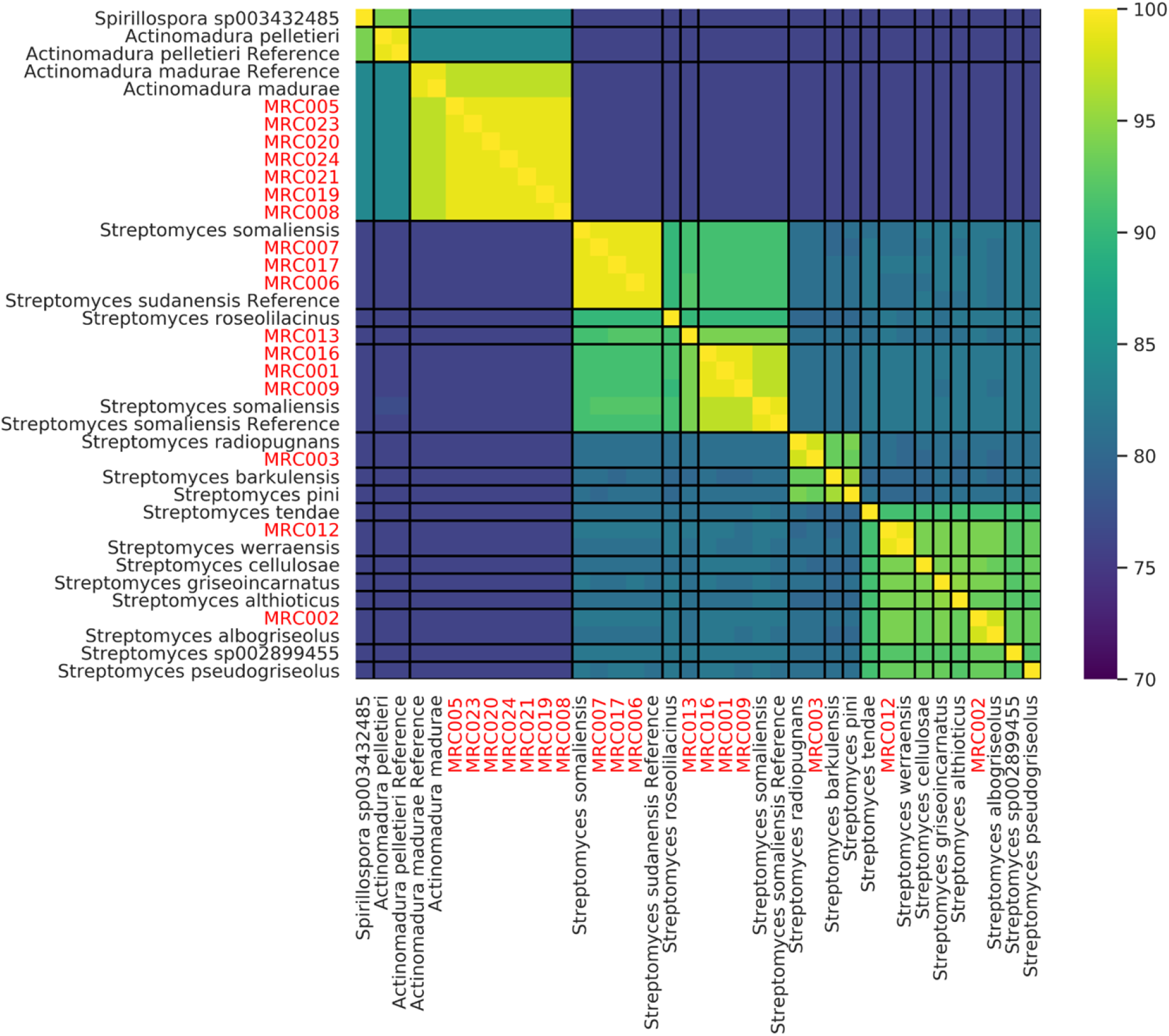
Pairwise comparison of Average Nucleotide Identity between the genomes of Isolates from the Mycetoma Research Centre (red) and their closest relatives in GTDB (all genomes from GTDB with an ANI >90 with any single isolate genome or the genome of type strains). The heatmap is ordered based on hierarchical clustering (Ward, Euclidian distance). Black lines delineate species boundaries based on ANI > 95%. The genomes of the reference species are from type strains.

**Fig 5.**
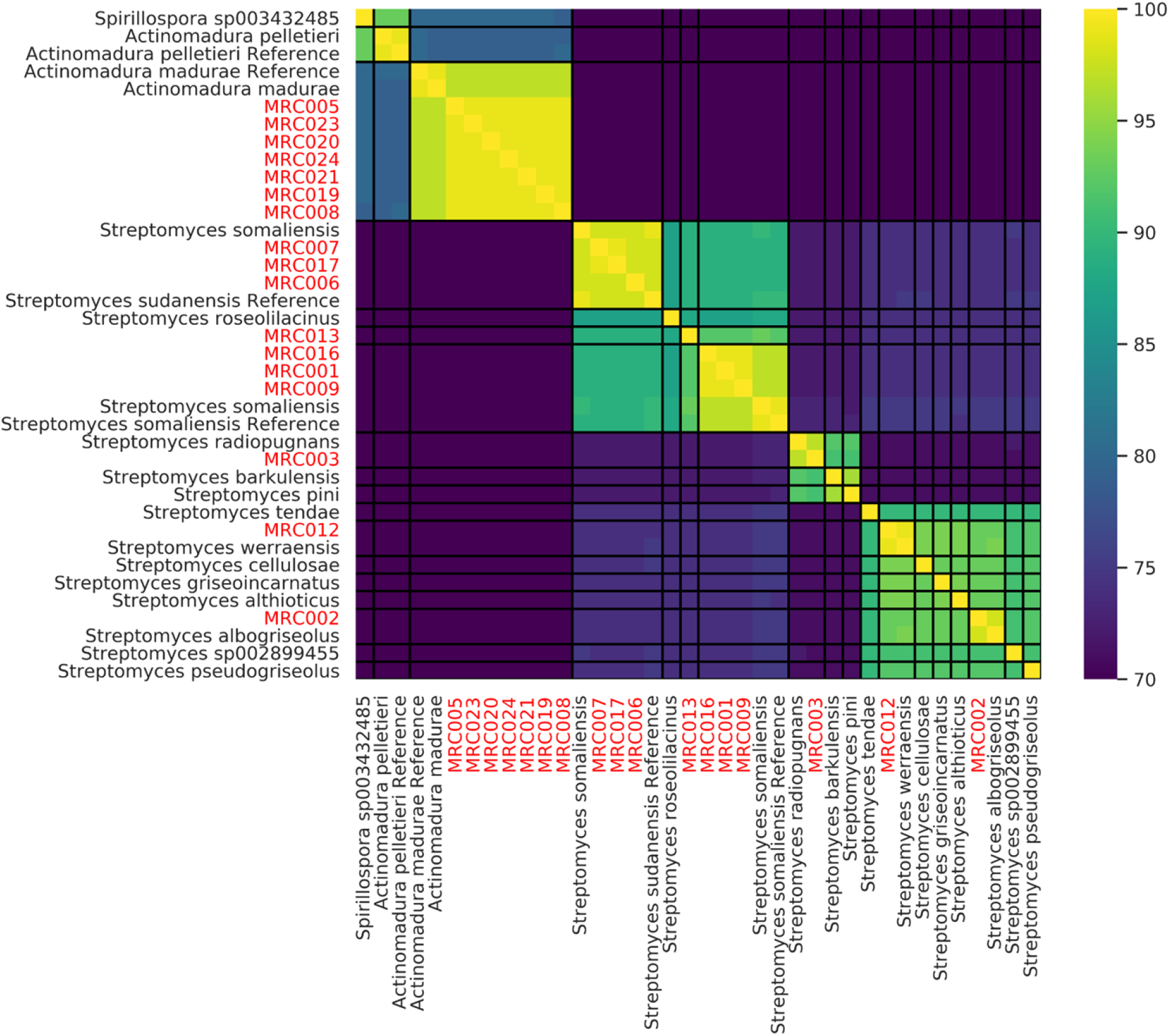
Pairwise comparison of Average Amino Acid Identity (AAI) between the genomes of Isolates from the Mycetoma Research Centre (red) and their closest relatives in GTDB (all genomes from GTDB with an ANI >90 with any single isolate genome or the genome of type strains). The heatmap order matches Fig 4. Black lines delineate species boundaries based on an AAI > 95%. The genomes of the reference species are from type strains.

**Table 2:** Summarising the pairwise ANI, AAI and estimated DDH values between isolates and their nearest relatives with sequenced genomes. *S. somaliensis* and *S. sundanensis* type strain genomes are compared to genomes in GTDB. Values that fall below defined thresholds for assigning strains to the same species are highlighted in grey.

| Genome 1 | Genome 2 | ANI (%) | AAI (%) | DDH (GGDH) |
| --- | --- | --- | --- | --- |
| MRC001 | *S. somaliensis* (GCF_012396115) | 97.7 | 97.6 | 80.5 |
| MRC002 | *S. albogriseolus* (GCA_014650475) | 98.4 | 98.2 | 85.1 |
| MRC003 | *S. radiopugnans* (GCF_900110735) | 98.6 | 98.0 | 83.5 |
| MRC005 | *A. madurae* (GCF_900115095) | 97.9 | 97.7 | 81.6 |
| MRC006 | *S. sudanensis* (GCF_000258595) | 99.0 | 98.8 | 89.7 |
| MRC007 | *S. sudanensis* (GCF_000258595) | 99.0 | 98.9 | 89.8 |
| MRC008 | *A. madurae* (GCF_900115095) | 97.9 | 97.6 | 81.7 |
| MRC009 | *S. somaliensis* (GCF_012396115) | 97.8 | 97.7 | 80.4 |
| MRC012 | *S. werraensis* (GCA_014656175) | 99.1 | 99.0 | 92.5 |
| MRC013 | *S. somaliensis* (GCF_012396115) | 94.2 | 93.0 | 52.9 |
| MRC016 | *S. somaliensis* (GCF_012396115) | 97.6 | 97.6 | 80.7 |
| MRC017 | *S. sudanensis* (GCF_000258595) | 99.1 | 98.9 | 89.9 |
| MRC019 | *A. madurae* (GCF_900115095) | 97.8 | 97.5 | 81.0 |
| MRC020 | *A. madurae* (GCF_900115095) | 97.8 | 97.5 | 81.3 |
| MRC021 | *A. madurae* (GCF_900115095) | 97.8 | 97.5 | 81.2 |
| MRC023 | *A. madurae* (GCF_900115095) | 97.8 | 97.5 | 81.1 |
| MRC024 | *A. madurae* (GCF_900115095) | 97.8 | 97.6 | 81.3 |

### *Streptomyces* strain MRC013, a new actinomycetoma pathogen

Interestingly, strain MRC013 was an outlier within the *S. somaliensis* clade, in both the 16S rRNA (Fig 1A) and the concatenated protein marker tree (Fig 3). In both trees it formed a strongly supported clade with the type strain of *S. somaliensis* and related isolates, but branched at the base of the group as an adjacent lineage. In the 16S rRNA gene tree, this isolate formed a strongly supported clade with a sequence previously isolated from a mycetoma patient (Sengupta, Goodfellow & Hamid NCBI access: EU544239.1; unpublished), adjacent to but excluding other isolates and the type strain rRNA. Whole genome comparisons support the annotation of MRC013 as a new species, as do low ANI, AAI and dDDH values of 94.2%, 93.0% and 52.9% with the genome of its closest relative, *S. somaliensis* (Table 2; Fig 4; Fig 5). Further, pairwise ANI and AAI comparisons of all isolate genomes revealed that MRC013 shares <95% ANI and <95% AAI with the other *Streptomyces* related genomes identified in the GTDB (Fig4; Fig 5). These data indicate that MRC013 is a member of a previously unrecognised *Streptomyces* species that is closely related to *S. somaliensis* and is a causal agent of human actinomycetoma. Based on its isolation from the region of West Kordofan, the name proposed for this new taxon is *Streptomyces kordofanensis.*

### The first reported isolation of members of three validly published *Streptomyces* species from actinomycetoma patients

The initial taxonomic assignment of three isolates from the genus *Streptomyces*, MRC002, MRC003 and MRC012, did not correspond to previously identified sources of human actinomycetoma. MRC002 16S rRNA was most similar to 16S rRNA from *Streptomyces atrovirens* in the EzBiocloud database, and phylogenetic analysis of 16S rRNA placed it in a lineage with this species and soil isolates from Sudan (Fig1B), though support for this clade was relatively low (88 ultrafast bootstrap support). In contrast, all whole-genome comparisons including GTDB-tk, the concatenated protein marker tree (Fig3), ANI, AAI and DDH similarities (Table 2) support the alternative annotation of MRC002 as *Streptomyces albogriseolus*, which was proposed for a neomycin producing strain originally isolated from soil [59].

Both 16S rRNA and all whole-genome and ANI and dDDH data (Fig 1D; Fig 3; Table 2) support the annotation of MRC003 as *S. radiopugnans.* The type strain of this species was isolated from a radiation-polluted soil in China and found to be markedly resistant to gamma radiation [60]. Halotolerant strains assigned to this species have been isolated from soil samples from Antarctica [61]. Similarly, corresponding data (Fig 1E, Fig 3, Table 2) support the assignment of MRC012 to *Streptomyces werraensis* [62], the type strain of which produces nonactin, a polyketide antibiotic. Additional strains assigned to this species have been isolated from soil and animal fecal samples in India [62,63]. It is particularly interesting that isolate MRCO12 forms a well defined clade (Fig 1E) with strains isolated from soil samples from Sudan and from an organism isolated from lesions of a donkey with withers (EU544241.1;), as mentioned above. It is evident from Figure 3 that this isolate forms a distinct branch that lies between the *S. somaliensis* and *S. sudanensis* lineages.

### Seven geographically separated *Actinomadura madurae* isolates with low genetic diversity

Most of the 16S rRNA *Actinomadura* sequences showed their highest similarities to *A. mexicana*, whilst in the 16S rRNA tree all samples formed a weakly supported clade that included the type strains of *A. madurae* and *A. darangshiensis* (Fig 2). Comparative analyses of whole genome sequences are effective in clarifying relationships between closely related species that are difficult to resolve using conventional taxonomic methods [64]. GTDB-tk assigned all isolates to the *A. madurae* lineage, a result supported by the concatenated multi-marker phylogeny of the isolates and their close relatives. All of the isolates formed a strongly supported clade with *A. madurae*, to the exclusion of *A. darangshiensis* (which is a strongly supported adjacent lineage to the *A. madurae* / isolate clade) and other characterised *Actinomadura* pathogens such as *A. latina, A. mexicana* and *A. pelletieri* (Fig 6). The ANI, AAI and DDH values support the classification of the isolates as *A. madurae* (Table 2). Despite the geographic separation of these isolates, MRC019 was from Yemen rather than Sudan, their genomes were highly similar, more so than for any other isolate lineage in the datasets (Fig 4; Fig 5).

**Fig 6.**
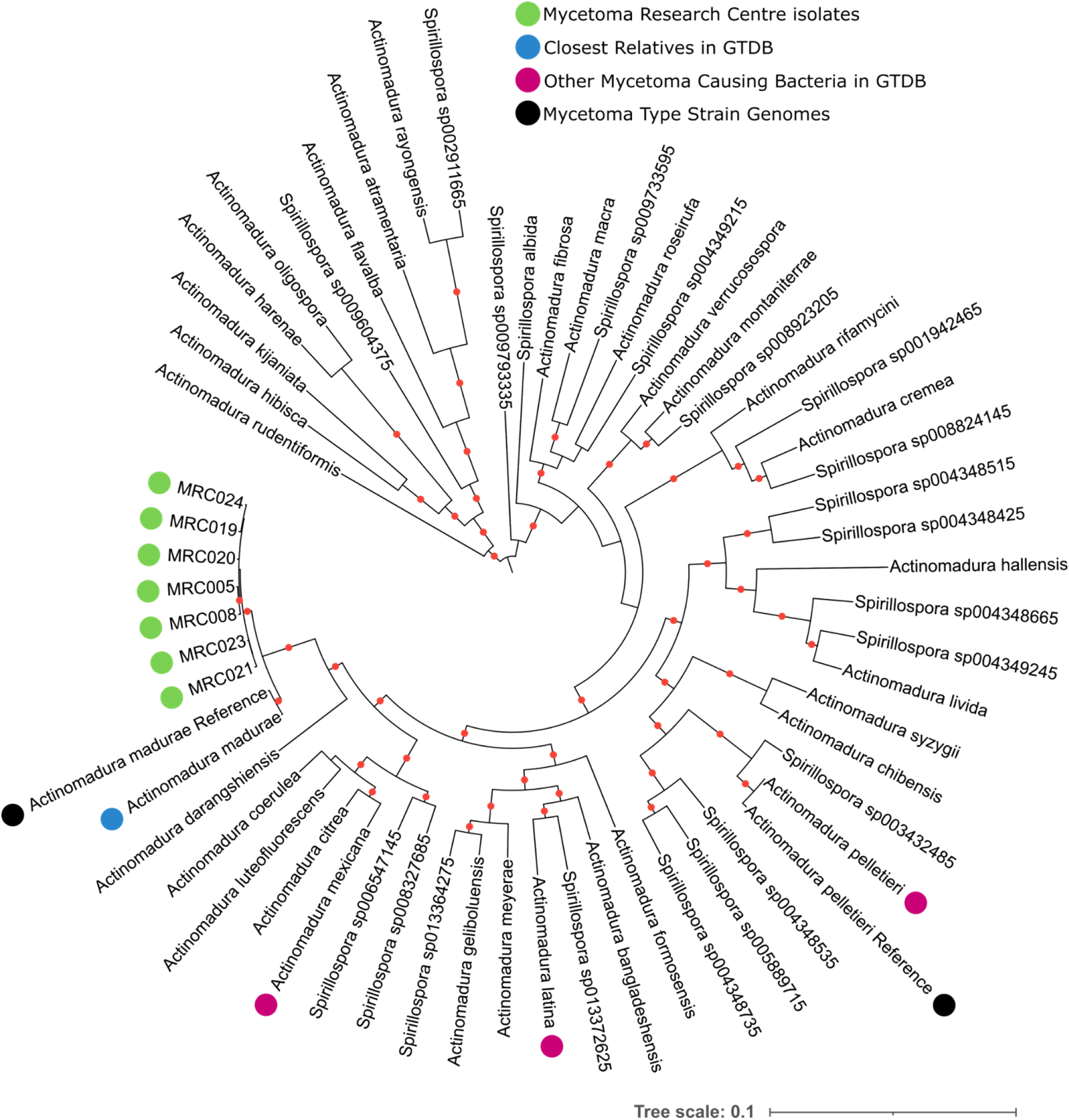
Tree of single copy orthologs that belong to the GTDB bac120 dataset from the genomes of *Actinomadura* related isolates from the Mycetoma Research Centre (green), their relatives according to ANI in GTDB (with the closest relatives indicated in blue) and genomes from type strains of species typically associated with actinomycetoma (black). Additionally, the type strain genome of any organism previously isolated from mycetoma patients and present in the GTDB is highlighted (pink). A red dot on branches indicates ultrafast bootstrap support >95. The tree was inferred in iqtree2 using the C20 model.

Basic statistics on the assembly of all genomes, alongside their final taxonomic assignments, are listed in Table 3. All of these genome sequences are publicly available under NCBI Bioproject PRJNA782605. It is interesting that the *S. somaliensis* and *S. sudanensis* strains have small genomes that range from 5.01 to 5.33 Mbp and 5.27 to 5.37 Mbp, respectively. The corresponding genome size for the putative type strain of *S. kordofanensis* is 5.33 Mbp. These comparatively low values for streptomycetes suggest that members of these taxa are undergoing an adaption from a saprophytic to a pathogenic lifestyle. It is also interesting that these strains have digital G+C contents that range from 73.93 to 74.18 %. In contrast, the *A. madurae* strains have much larger genomes and lower G+C contents, as shown in Table 3.

**Table 3:**
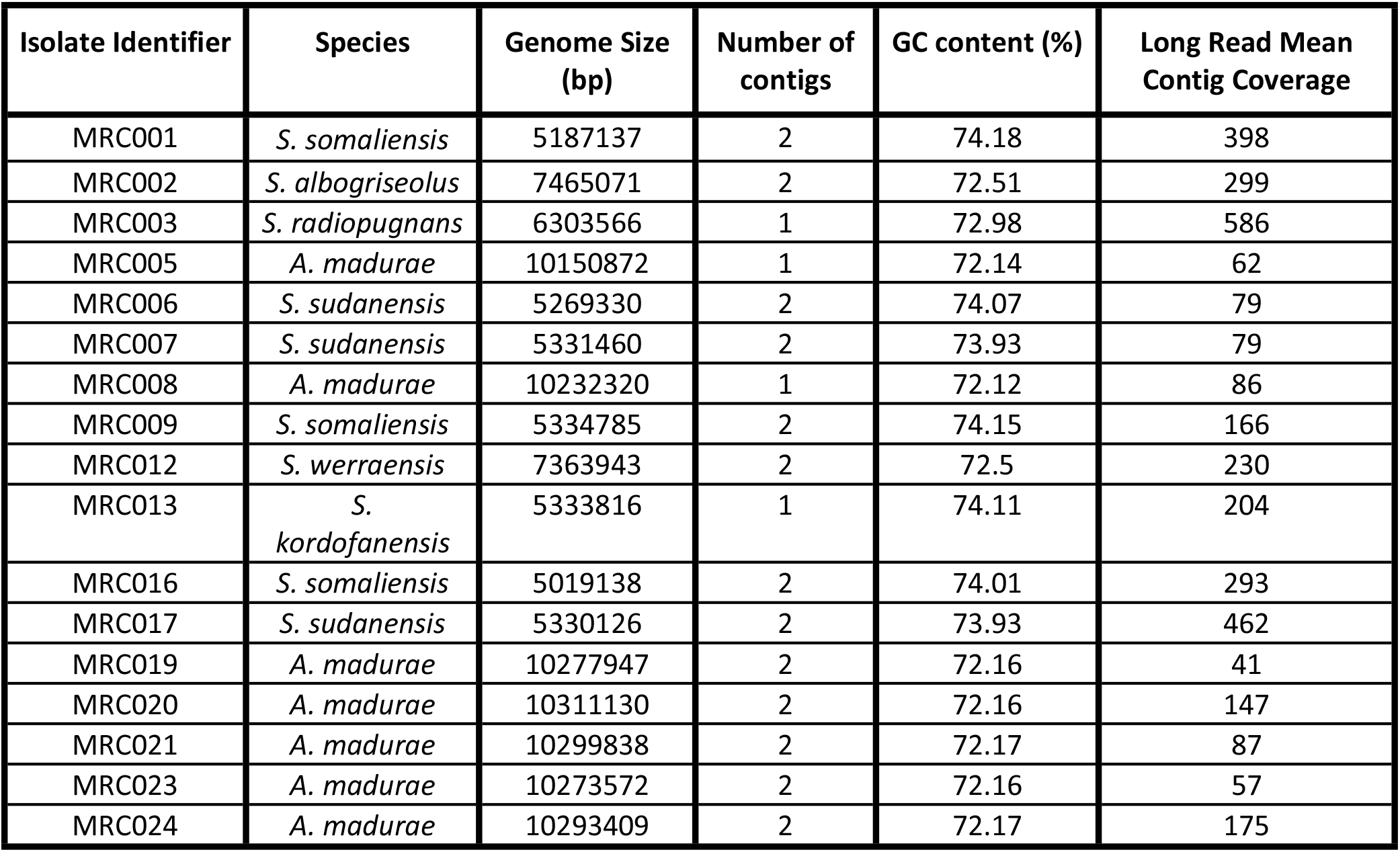
Assembly statistics for all genomes sequenced in this project. The genomes from these assemblies are available under NCBI Bioproject PRJNA782605.

### Discrepancies between phenotypic and molecular based classifications

The initial taxonomic classification of 8 of the 15 isolates based on histopathology or cytology was not supported by the taxonomic data (Table 4). Taxonomic classifications based on limited numbers of phenotypic traits are of limited reliability, so discrepancies between these original classifications and those from the genomic analyses are not surprising. These discrepancies are noteworthy due to their potential impact on treatment options. In two cases (MRC002 and MRC005), the initial classification as fungal (eumycetoma) rather than bacterial (actinomycetoma) pathogens led to lengthy treatment with antifungal agents.

**Table 4.** Initial clinical classification of isolates compared to the molecular taxonomic assignments.

| Isolate Metadata |  |  |  | Classification |  |  |
| --- | --- | --- | --- | --- | --- | --- |
| Strain number | Duration (years) | Lesion Size (cm) | Locality | Histopathology | Cytology | Molecular |
| MRC001 | 1 | >10 | Khartoum | *S. somaliensis* | ND | *S. somaliensis* |
| MRC002 | 7 | > 10 | White Nile | *S. somaliensis* | *Madurella mycetomatis* | *S. albogriseolus* |
| MRC003 | 1 | NK | North Kordofan | *S. somaliensis* | *S. somaliensis* | *S. radiopugnans* |
| MRC005 | 1 | 5-10 | Khartoum | ND | *M. mycetomatis* | *A. madurae* |
| MRC006 | 1 | 5-10 | South Darfour | ND | *A. madurae* | *S. sudanensis* |
| MRC007 | 4 | < 5 | White Nile | *S. somaliensis* | ND | *S. sudanensis* |
| MRC008 | 1 | > 10 | South Kordofan | ND | *S. somaliensis* | *A. madurae* |
| MRC009 | 30 | > 10 | White Nile | ND | *S. somaliensis* | *S. somaliensis* |
| MRC012 | 20 | > 10 | Kassala | ND | *A. madurae* | *S. werraensis* |
| MRC013 | 3 | 5-10 | West Kordofan | *A. madurae* | *A. madurae* | *S. kordofanensis* |
| MRC016 | 1 | NK | White Nile | *S. somaliensis* | *S. somaliensis* | *S. somaliensis* |
| MRC017 | 1.5 | > 10 | North Kordofan | *A. madurae* | ND | *S. sudanensis* |
| MRC019 | 8 | > 10 | Yemen | ND | *A. pelletieri* | *A. madurae* |
| MRC020 | 8 | > 10 | White Nile | *S. somaliensis* | ND | *A. madurae* |
| MRC021 | 4 | > 10 | North Kordofan | *A. madurae* | ND | *A. madurae* |
| MRC023 | 1 | > 10 | North Kordofan | ND | *A. madurae* | *A. madurae* |
| MRC024 | 4 | 5-10 | North Kordofan | ND | *A. madurae* | *A. madurae* |

NK, lesion size not known. ND, not determined.

### In vivo and in vitro antibiotic resistance profiling

The first-line treatment of actinomycetoma is a combination of sulfamethoxazole/trimethoprim and amoxicillin/clavulanic acid [35]. To verify the sensitivity of our isolates to these antibiotics and explore potential alternative treatments, we tested each isolate against a range of commonly used antibiotics under laboratory conditions. To our surprise, most isolates were fully resistant to sulfamethoxazole/trimethoprim (Fig 7). In general, the *S. somaliensis* and *S. sudanenis* strains were more susceptible to the tested antibiotics. The β-lactamase inhibitor clavulanic acid enhanced the sensitivity of most strains to amoxicillin, suggesting that these strains produce a β-lactamase. The *A. madurae* isolates were more resistant to all tested antibiotics, apart from amikacin.

**Fig 7.**
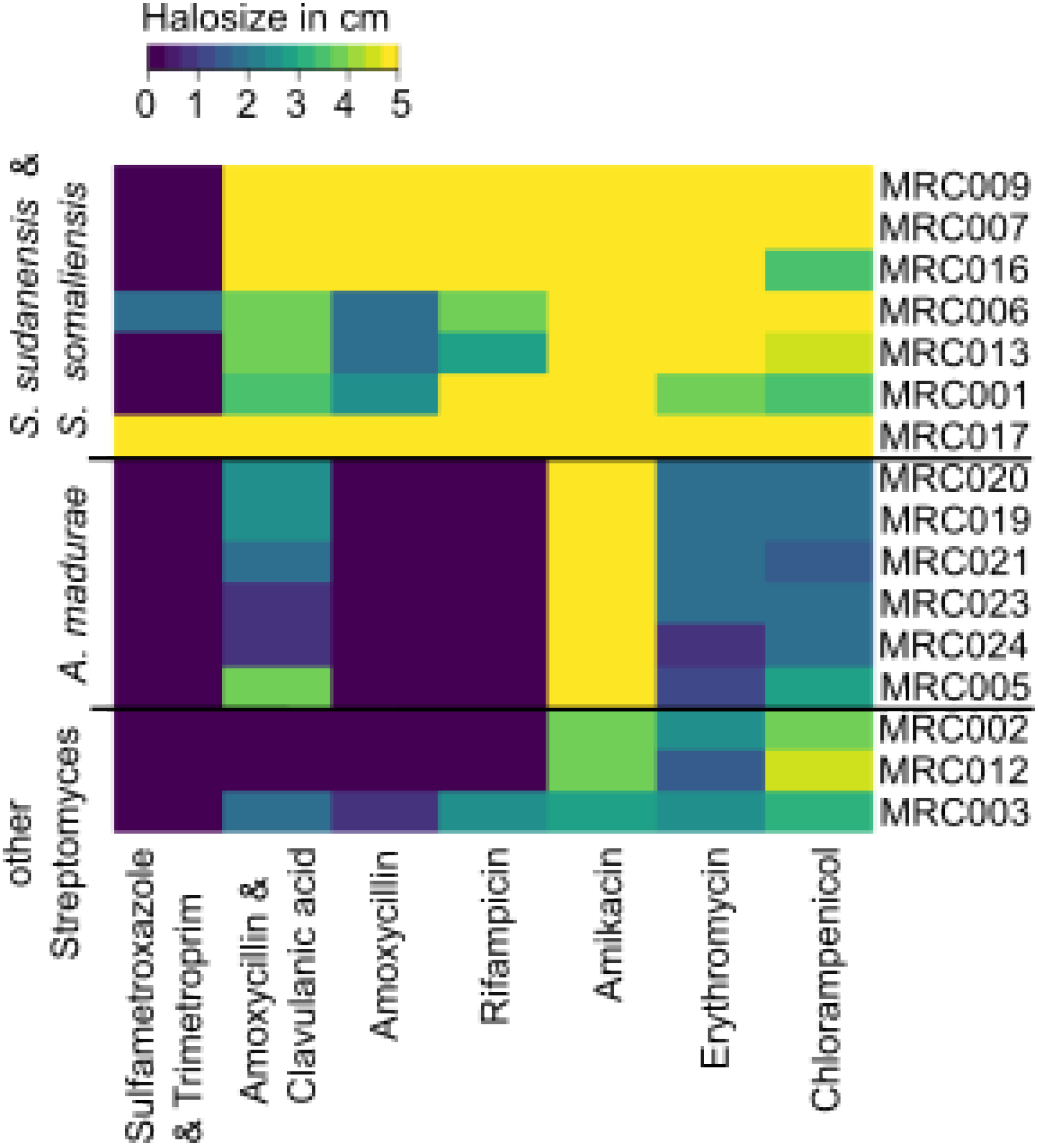
Comparative antibiotic susceptibility profiles of actinomycetoma isolates based on a disk diffusion assay. Halo size was recorded in cm. Yellow most susceptible. Blue non-susceptible. The isolates were grouped using hierarchical clustering based on their resistance profiles using the default parameters of heatmap2 in R.

In the light of these findings, we scanned the whole genome sequences for known antibiotic resistance determinants using AMRfinderPLUS (SI Fig S2). This indicated the presence of rifampicin resistance genes in *A. madurae*, a result in agreement with the observed resistance profiles. The *A. madurae* strains also had 2 genes annotated as β-lactamases, while the *S. somaliensis* and *S. sudanensis* strains only had one putatively conferring the higher level of resistance to β-lactam antibiotics observed *in vivo*. These results suggest that whole-genome sequencing can be used to predict antibiotic susceptibility and guide treatment. Alternatively, the distribution of common resistance genes in the genome sequences could be used to generate rapid diagnostic PCR-based tests.

## Discussion

The taxonomic diversity of bacteria that can cause actinomycetoma contributes to the difficulty in accurately diagnosing and treating the disease [26,28]. In this study, we compared the currently used phenotypic-based methods for diagnosing mycetoma, such as histopathology, cytology, and micromorphological appearance, with whole-genome sequencing data. While all strains isolated belonged to the phylum *Actinobacteria*, as predicted by at least one of the classical phenotypic methods, the molecular analysis highlighted several cases in which the causative agent of infection was misdiagnosed. It is also significant that the molecular sequence and genotypic data confirmed the species status of *A. latina [18,19]* and *S. sudanensis [23]*. Furthermore, the identification of four *Streptomyces* species with no previous association with human mycetoma in this relatively small dataset, including isolate MRC013 which is the first sequenced representative of the proposed new taxon *S. kordofanensis*, expands the range of *Streptomyces* species known to be associated with the disease. Further work with broader sample sizes are needed to establish how important these new species are in the overall worldwide disease burden, and whether more species capable of causing actinomycetoma remain to be discovered, as seems likely [19,65].

The identification of isolates of *S. sudanensis* and *S. warraenensis* with similarity to samples previously isolated from donkey withers by Sengupta, Goodfellow and Hamid (unpublished) suggests that some actinomycetoma causing strains may have the potential to infect multiple host species [42]. Epidemic outbreaks of eumycetoma (*Sporothrix brasiliensis*) have previously been linked with zoonotic transmission from cats to humans [66], and the identification of multi-host actinomycetoma causing strains also raises zoonosis as a possible route for transmission of the bacterial disease.

Actinomycetes are best known for their ability to produce antibiotics and often encompass a multitude of antibiotic resistance genes [67]. To test whether the accurate diagnosis of the causative agent of actinomycetoma infections could positively impact treatment outcomes for the disease, we investigated the resistance of isolates to a range of commonly used antibiotics. In general, all strains sequenced showed elevated levels of laboratory resistance to some antibiotics. It is possible that even higher levels of resistance operate in physiological contexts, for example, within grains. The current first-line treatments for actinomycetoma are long-term administration with sulfamethoxazole/trimethoprim and amoxicillin/clavulanic acid. It was surprising that almost all our isolates had high-level resistance to sulfamethoxazole/trimethoprim under laboratory conditions. In addition, most bacteria were highly resistant to amoxicillin. *A. madurae* and two of the previously unidentified isolates, MRC002 and MRC0012, were particularly resistant, highlighting the potential importance of accurate pathogen identification for treatment choices. Nevertheless, these drugs appear to be effective in a clinical setting, so more work is needed to understand the link between the choice of therapy and the clinical outcome. The activity of amoxicillin could be rescued to a degree by the addition of the β-lactamase inhibitor clavulanic acid, as commonly used in clinical settings. While alternative drugs tested here, such as amikacin, seemed to be more effective, it is important to note that they have considerable toxic side effects [34], hence the benefit may not outweigh the risk for the patient.

In conclusion, the present study shows that the current diagnostic tests used to identify the causative agents of mycetoma have serious limitations. Two of the patients were diagnosed with eumycetoma and given lengthy anti-fungal treatments with common negative side effects and complications, but were thereafter shown to have actinomycetoma. Given the observed differences in antimicrobial resistance and misdiagnosis of infections, a wider study of actinomycetoma pathogens from around the world is urgently needed, in combination with the development of point of care rapid molecular diagnostics. Furthermore, antimicrobial resistance profile testing should be available at mycetoma clinics to avoid giving patients inappropriate antibiotics, leading to increased morbidity and drug resistance.

The status of *S. kordofanensis* as a new species within the genus *Streptomyces* is mainly based on phylogenomic analyses of closely related reference strains, notably the type strains of *S. somaliensis* and *S. sudanensis*.

### Description of *Streptomyces kordofanensis* sp. nov. *Streptomyces kordofanensis* (kor.do.fan.en’ sis. N. L. masc. adj. *kordofanensis*, belonging to Kordofan, the source of the isolate)

Aerobic, Gram-stain positive actinomycete which forms an extensively branched pale/creamy yellow colour substrate mycelium on Tryptic Soy Agar. Neither aerial hyphae nor diffusible pigments are formed on this medium. Grows from pH6 to pH9, optimally at pH7 from 30 to 37C. Colonies are convex and have filamentous margins. Susceptible to amoxicillin (with the effect enhanced by the addition of clavulinic acid), rifampacin, amikacin, erythromycin and chloramphenicol., but resistant to sulfamethoxazole and trimetroprim. The digital GC content is 74.11% and the genome length 5.33 Mb, assembled into a single contig.

The type and only strain, MRC013T (= DSMXXXX = NCIMBXXX) was isolated from granulomatous material of mycetoma lesions of a patient in the Sudan. The accession number of the 16S rRNA gene sequence of the isolate is XXXX And that of the genome sequence is SAMN23388489.

## Acknowledgement

We are grateful to Aharon Oren (the Hebrew University of Jerusalem) for checking the species epithet of the novel *Streptomyces* isolate.

## Author Contributions

Andrew Keith Watson: Data Curation, Formal Analysis, Investigation, Software, Visualization, Writing – Original Draft Preparation, Writing – Review & Editing

Bernhard Kepplinger: Conceptualization, Data Curation, Formal Analysis, Investigation, Methodology, Visualization, Writing – Original Draft Preparation, Writing – Review & Editing

Sahar Mubarak Bakhiet: Data Curation, Investigation, Methodology, Resources

Nagwa Adam Mhmoud: Investigation, Methodology, Resources.

Michael Goodfellow: Conceptualization, Writing – Original Draft Preparation, Writing – Review & Editing.

Ahmed Hassan Fahal: Conceptualization, Writing – Original Draft Preparation, Writing – Review & Editing

Jeff Errington: Conceptualization, Funding Acquisition, Project Administration, Supervision, Writing – Original Draft Preparation, Writing – Review & Editing

